# GATA2 Mediates Macrophage Proliferation During Atherosclerosis

**DOI:** 10.1101/2025.02.18.638845

**Authors:** Amena Aktar, Angela M Vrieze, Kiera Telesnicki, Minhyuk Mun, Kasia Wodz, Charles Yin, Paisley Cox-Duvall, David Nagpal, Bryan Heit

## Abstract

Atherosclerosis is fueled by the buildup of lipid-laden macrophages within the vascular intima. These macrophages are derived from monocytes that are recruited from the circulation into the developing lesion, where they proliferate and differentiate into macrophages, with proliferation producing most of the macrophages in the resulting lesions. However, the signals and transcriptional events driving the proliferation of atherosclerotic plaque macrophages remain poorly understood. Analysis of human plaque spanning a range of disease severity identified a subpopulation of macrophages that expressed the hematopoietic transcription factor GATA2. These GATA2-expressing macrophages had a transcriptional profile that was intermediary between monocytes and mature macrophages, and selectively upregulated genes associated with proliferation and apoptosis. The expression of GATA2 was concomitant with plaque macrophage proliferation at all stages of disease, with over 90% of proliferating macrophages expressing GATA2. GATA2 was upregulated in macrophages following exposure to oxLDL, with GATA2 expression being necessary and sufficient for the proliferation of these macrophages. In these cells, GATA2 mediates proliferation by upregulating expression of the proto-oncogene MYB, while simultaneously decreasing sensitivity to apoptosis induced by the unfolded protein response. Together, these data identify GATA2 as a transcription factor upregulated by atherogenic stimuli that functions as the primary mediator of macrophage proliferation in atherosclerotic plaque.

## Introduction

Macrophages are the primary cell type involved in the development of atherosclerosis, where they serve both protective and atherogenic roles [1]. In early disease, oxidized Low-Density Lipoprotein (oxLDL) and other modified forms of LDL are deposited within the intima of the vasculature. These deposits are engulfed by macrophages via the scavenger receptor CD36, which then export the cholesterol to High-Density Lipoprotein (HDL), thereby removing the deposits from the vascular intima and returning the material to the circulation [2,3]. Excess uptake of oxLDL can overwhelm this system, causing macrophages to accumulate cholesterol and lipids in intracellular droplets, forming foam cells [4]. Foam cells are highly inflammatory and promote the ingress of monocytes which differentiate into additional macrophages. These foam cells eventually die due to activation of the unfolded protein response that is induced by the intracellular stresses created by the accumulated lipids and sterols [5]. Normally, other macrophages would clear these dying cells through efferocytosis, but for unclear reasons this activity is lost early in disease [6,7]. The loss of efferocytosis results in the accumulation of apoptotic foam cells, which uncleared, undergo secondary necrosis and spill their accumulated lipid and sterol content back into the extracellular space [8]. This accumulation of dead macrophages and the cell-free lipids and sterols released from these cells forms the necrotic core that typifies atherosclerotic lesions. In early disease most plaque macrophages are derived from monocytes recruited from the vasculature, but as disease progresses the growth of the plaque macrophage population becomes increasingly dependent on macrophage proliferation and less dependent on monocyte influx [9].

In mouse models, plaque macrophages undergo turnover in just four weeks, with local proliferation accounting for nearly 90% of macrophage replenishment in advanced plaques [9]. Consistent with this, targeting macrophage proliferation is sufficient to reduce plaque inflammation and promote plaque remodeling to a less pathological state [10]. In fact, some studies suggest that the anti-proliferative impact of statins may account for the majority of their impact on plaque regression, exceeding the contribution of statins on plaque inflammation and lowering cholesterol levels [11]. Plaque macrophage proliferation is initiated by oxLDL-induced scavenger receptor signaling, which then activates a downstream signalling cascade dependent on AMPK, PI3K, p38 MAPK, and ERK [12–15]. This cascade is further amplified by oxLDL-induced autocrine signaling via M-CSF and GM-CSF [16,17]. But while the ligands and upstream signaling molecules mediating this macrophage proliferation are established, the specific transcriptional targets and cell cycle regulatory pathways involved in this process remain largely unknown.

We recently identified a population of GATA2-expressing macrophages in small, early-stage plaques found in the intima of aortic biopsies recovered during coronary artery bypass graft surgery [18]. This population of macrophages accounted for a large portion of the macrophages present in these lesions, and *in vitro* exhibited several atherogenic phenotypes including impaired cholesterol export and defects in apoptotic cell removal and processing [18]. Finding GATA2 expression in these cells was unexpected, as previous literature suggested that GATA2 expression in myeloid cells is limited to the bone marrow (reviewed in [19]). Within the bone marrow, GATA2 is required for the proliferation of hematopoietic stem cells and myeloid precursor cells prior to their commitment to terminal differentiation [20]. Indeed, the only other context in which myeloid cells regularly express GATA2 outside of the bone marrow niche is in myeloid leukemia and myelodysplastic syndrome, where GATA2 expression promotes cell cycle progression and malignancy [19]. GATA2 is a member of the GATA family of transcription factors, and binds to the DNA motif “GATAA” via a tandem zinc finger motif [21,22]. GATA2 can function as a pioneering transcription factor, opening heterochromatin to form euchromatin that is accessible to other transcription factors and RNA polymerase [23]. In bone marrow, GATA2 expression is required for the self-renewal of hematopoietic stem cells and hematopoietic progenitor cells. GATA2 expression peaks when these cells commit to the myeloid lineage, with GATA2 expression then decreasing during terminal differentiation [24]. GATA2 deficiency is lethal due to the loss of hematopoietic stem cells in early embryogenesis [25], with mutations that reduce GATA2 activity leading to the development of conditions such as MonoMac syndrome— characterized by monocytopenia, NK- and B-lymphocytopenia, and a susceptibility to the development of myelodysplasia and acute myeloid leukemia [26–28].

GATA2 expression oscillates during the cell cycle, with the highest expression occurring during S-phase, with this cyclical expression pattern thought to regulate the proliferation of hematopoietic stem cells [29,30]. During G1 phase, GATA2 expression is suppressed by CDK4 [31]. Loss of this suppression increases GATA2 expression, where via inducing the expression of MYB, GATA2 mediates the G1/S transition [32]. Once in S-phase, GATA2 induces expression of cyclin D1 and cyclin E1, which initiates a positive feedback loop that sustains GATA2 expression through G2 phase [31]. The high levels of these cyclins and GATA2 then mediates passage through the G2/M checkpoint, thereby completing the proliferative cycle. Given the key role of GATA2 in the cell cycle, and its expression in plaque macrophages, we used human atheroma samples and human macrophage cell lines to test the hypothesis that GATA2 expression mediates the proliferation of plaque macrophages.

## Results

### GATA2+ MAcrophages Are Monocytes Arrested During Macrophage Differentiation

We previously determined that a large portion of macrophages present in early-stage aortic atheromas express GATA2, but the ontology and role of these GATA2+ macrophages in atherosclerosis remains unclear [18]. Using aortic biopsies collected during coronary artery bypass graft surgery, we determined that the density of macrophages in these nascent plaques varied greatly, with the fraction of macrophages expressing GATA2 positively correlated with macrophage density (**Figure 1A-B**). To better understand the phenotype of these GATA2+ macrophages, we purified these cells via laser microdissection and compared their transcriptome to that of monocytes and monocyte-derived macrophages (moMacs) from healthy donors. Principal component analysis determined that the gene expression profile of GATA2+ macrophages falls mid-way between monocytes and moMacs, suggesting that the GATA2+ macrophages may be monocytes arrested in their differentiation into plaque macrophages or are macrophages which have partially de-differentiated into an immature state (**Figure 1C**). A detailed transcriptomic analysis of these cells identified over 5,100 genes that were differentially regulated between monocytes, GATA2+ macrophages, and moMacs. Consistent with the principal component analysis, GATA2+ macrophages exhibited partial downregulation of monocyte-expressed genes (Cluster 6, **Figure 1D**), and partial upregulation of macrophage-expressed genes (Clusters 1, 3-5, **Figure 1D**). In addition to this intermediary gene expression profile, GATA2+ macrophages also upregulated a small number of genes that were minimally expressed in both monocytes and moMacs (Cluster 2, **Figure 1D**). Gene ontology analysis of Cluster 2 identified an enrichment of genes involved in cell cycle and apoptosis (**Figures 1E, S1**). Consistent with this, several genes promoting cell proliferation and negatively regulating apoptosis were selectively upregulated in GATA2+ macrophages, including GATA2 (**Figure 1F-G**). All three cell types lacked expression of the yolk-sac erythro-myeloid progenitor marker Runx1, and exhibited expression of Kit, Tie2/TEK, and Flt3, consistent with an adult hematopoietic ontology (**Figure 1H**)[33]. However, GATA2+ macrophages only formed when monocytes from atherosclerosis patients were differentiated into macrophages, whereas moMacs differentiated from the monocytes of healthy controls lacked GATA2 expression (**Figure 1I**), suggesting that the upregulation of GATA2 during monocyte-to-macrophage differentiation is specific to atherogenic conditions, and is not a normal part of monocyte-to-macrophage differentiation.

**Figure 1:**
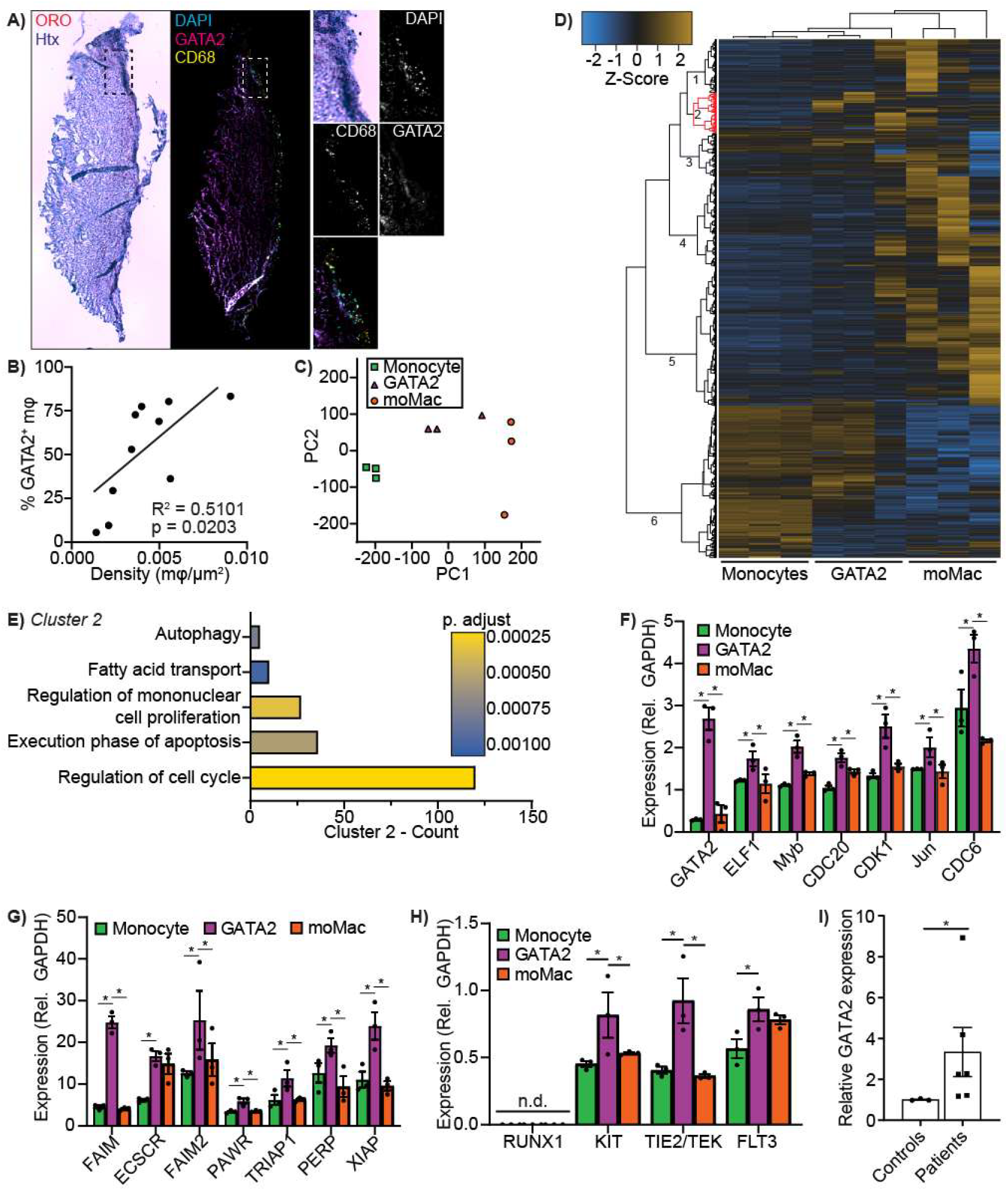
GATA2+ Macrophages Exhibit a Transcriptional Profile Intermediary to Monocytes and Monocyte-Derived Macrophages. **A)** Micrographs of serial sections of an aortic punch biopsy, stained for lipids (Oil-Red-O, ORO) and cells (Hematoxylin, Htx), or fluorescently stained for DNA (DAPI, cyan), GATA2 (magenta), and macrophages (CD68, yellow). Insert shows an early-stage plaque identified as a region containing foci of oil-red-o staining and increased cellularity. Scale bar is 100 μm. **B)** Correlation between the frequency of GATA2+ macrophages (m ϕ) and macrophage density within early-stage atherosclerotic lesions, analyzed by a Spearman’s correlation coefficient. **C)** Principal component analysis of the transcriptome of monocytes, GATA2+ macrophages (GATA2), and monocyte-derived macrophages (moMac). **D)** Transcriptome heatmap comparing monocytes, GATA2+ macrophages (GATA2), and monocyte-derived macrophages (moMac). Six clusters of genes under similar regulation were identified (numbered 1-6), with cluster 2 (red) containing genes selectively upregulated in GATA2+ macrophages. **E)** Gene ontology (GO) analysis of Cluster 2. Count is the number of genes in cluster 2 assigned to the corresponding GO annotation. **F-G)** Top differentially expressed genes in cluster 2 that are assigned to GO annotations for cell proliferation (F) and apoptosis (G). **H)** Expression of macrophage markers of embryonic (Runx1) versus hematopoietic (Kit, Tie2/TEK and Flt3) ontology in monocytes, GATA2+ macrophages (GATA2), and monocyte-derived macrophages (moMac). n.d. = not detected. **I)** RT-PCR quantification of GATA2 expression in monocyte-derived macrophages. GATA2 expression is normalized to GAPDH and expressed as fold-expression relative to healthy controls. * p < 0.05 between indicated groups, Kruskal-Wallis with Dunn correction (F-H) or Mann-Whitney U test (H).

### Monocytic gata2 expression Does Not Drive Macrophage gata2 expression

Given that GATA2+ macrophages were found in early-stage atheromas of all tested patients, where GATA2 may function to arrest monocyte-to-macrophage differentiation, we used flow cytometry to determine whether GATA2 was expressed in the monocytes of the same patients (**Figure 2A**). GATA2 was minimally expressed in most healthy controls and patient monocytes, with a small portion of individuals exhibiting high levels of monocytic GATA2 expression (**Figure 2B**). No correlation between the expression level of GATA2 in patient monocytes and the frequency of GATA2+ macrophages in the patients’ atherosclerotic plaques was observed (**Figure 2C**). While there was no correlation between monocyte and macrophage GATA2 expression, the differing levels of GATA2 expression in patient monocytes may correlate with disease severity. To assess this possibility, we divided patients into GATA2-high (GATA2 MFI > 2 S.D. above healthy controls) and GATA2-low populations, then compared the expression of several monocyte-associated markers that were expressed at high levels in GATA2+ macrophages between these populations (**Figure 1D**). While atherosclerosis patients had an overall higher expression of the chemokine receptor CCR2, and of the α_4_ and β_2_ integrins than controls, there were no differences in the expression of these genes between GATA2-high and GATA2-low patient monocytes (**Figure 2D, E**). Consistent with these observations, no differences in the recruitment of GATA2-high versus -low patient monocytes to TNFα-stimulated human aortic endothelium were observed in a parallel-plate flow chamber assay of monocyte recruitment (**Figure S2**).

**Figure 2:**
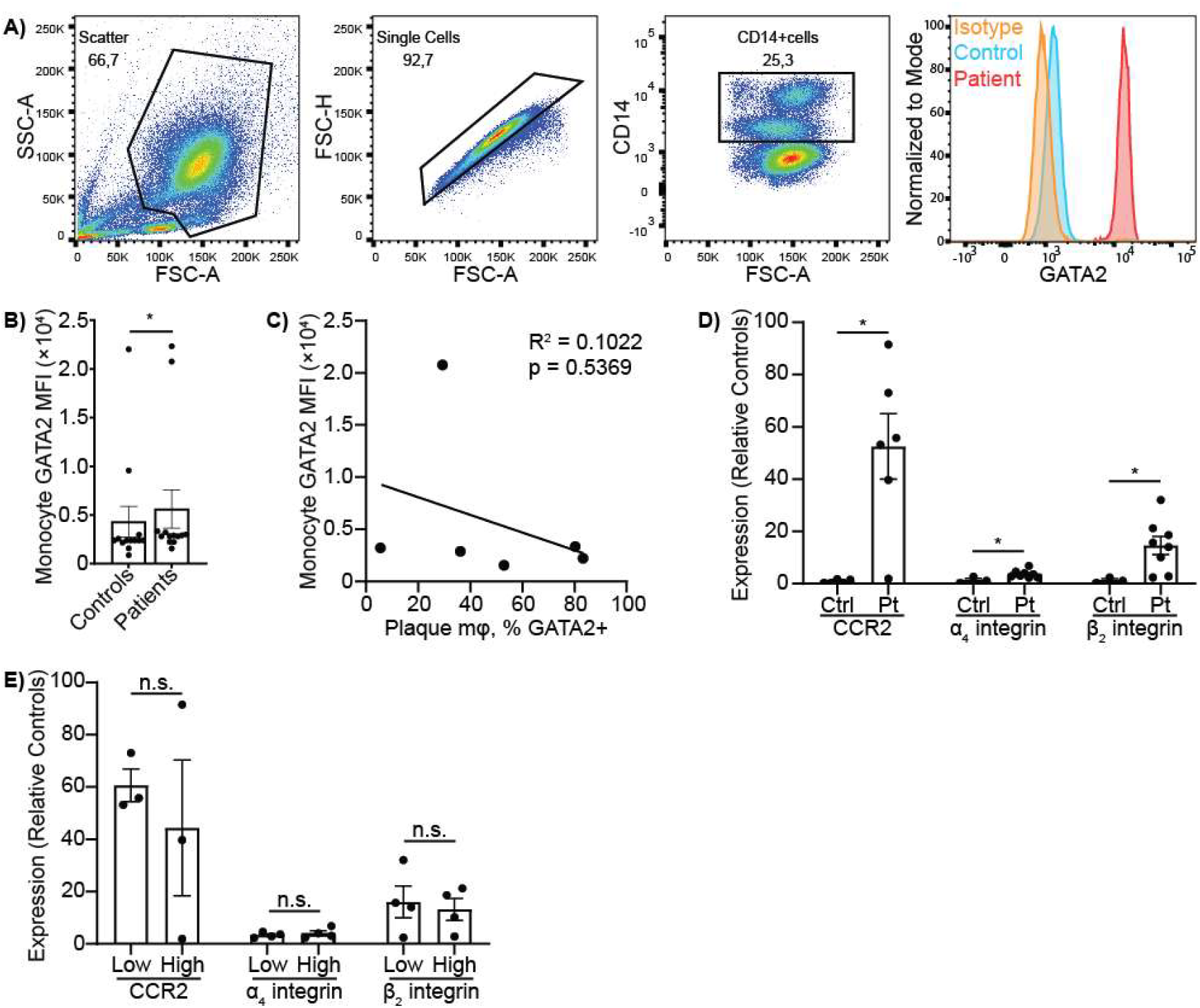
GATA2 is expressed in the monocytes of a portion of atherosclerosis patients. Flow cytometry was used to quantify the expression of GATA2 and other monocyte markers in peripheral blood mononuclear cells isolated from heathy controls and atherosclerosis patients undergoing coronary artery bypass graft surgery. **A)** Flow cytometry gating strategy for quantifying GATA2 expression in CD14+ monocytes. **B)** Expression of GATA2 in monocytes from healthy controls and atherosclerosis patients. **C)** There is no correlation between the mean fluorescence intensity (MFI) of GATA2 in patient monocytes versus the fraction of macrophages expressing GATA2 in early-stage plaques from aortic punch biopsies taken during coronary artery bypass graft surgery. **D**,**E)** Expression of CCR2, α_4_ integrin, and β_2_ integrin in the monocytes of healthy control versus atherosclerosis patients (D) and in patients with high (MFI >2 S.D. compared to healthy controls, “High”) versus low monocytic GATA2 expression. n = minimum of 3, * = p < 0.05, n.s. = p > 0.05 between indicated groups, Mann-Whitney U test (B,D,E), or Spearman’s correlation coefficient (C).

To further explore potential links between monocyte GATA2 expression and markers of disease severity, we compared the cytokine, lipid, and sterol profiles of patients with GATA2-low versus -high monocytes. Luminex analysis of plasma cytokines identified minimal differences between controls and both GATA2-low or GATA2-high patients (**Figure 3A**). Only two cytokines were significantly different between healthy controls and GATA2-high patients, with GATA2-high patients exhibiting an ∼2-fold increase in serum PDGF-AB/BB and a <90% decrease in serum IL-23 (**Figure 3B-C**). GATA2 was modestly upregulated when monocytes were differentiated into moMacs in the presence of 12 ng/mL PDGF-AB/BB, but this increase was not observed at other PDGF-AB/BB concentrations (**Figure 3D**). Patients with GATA2-high monocytes had a modest increase in non-HDL cholesterol but were otherwise similar to GATA2-low patients across other lipid measures (**Figure 3E**). GATA2 expression was not induced in moMacs cultured with LDL (**Figure 3F**). These results indicate that plaque-derived stimuli, not circulating factors, drive GATA2 expression in early atheromas.

**Figure 3:**
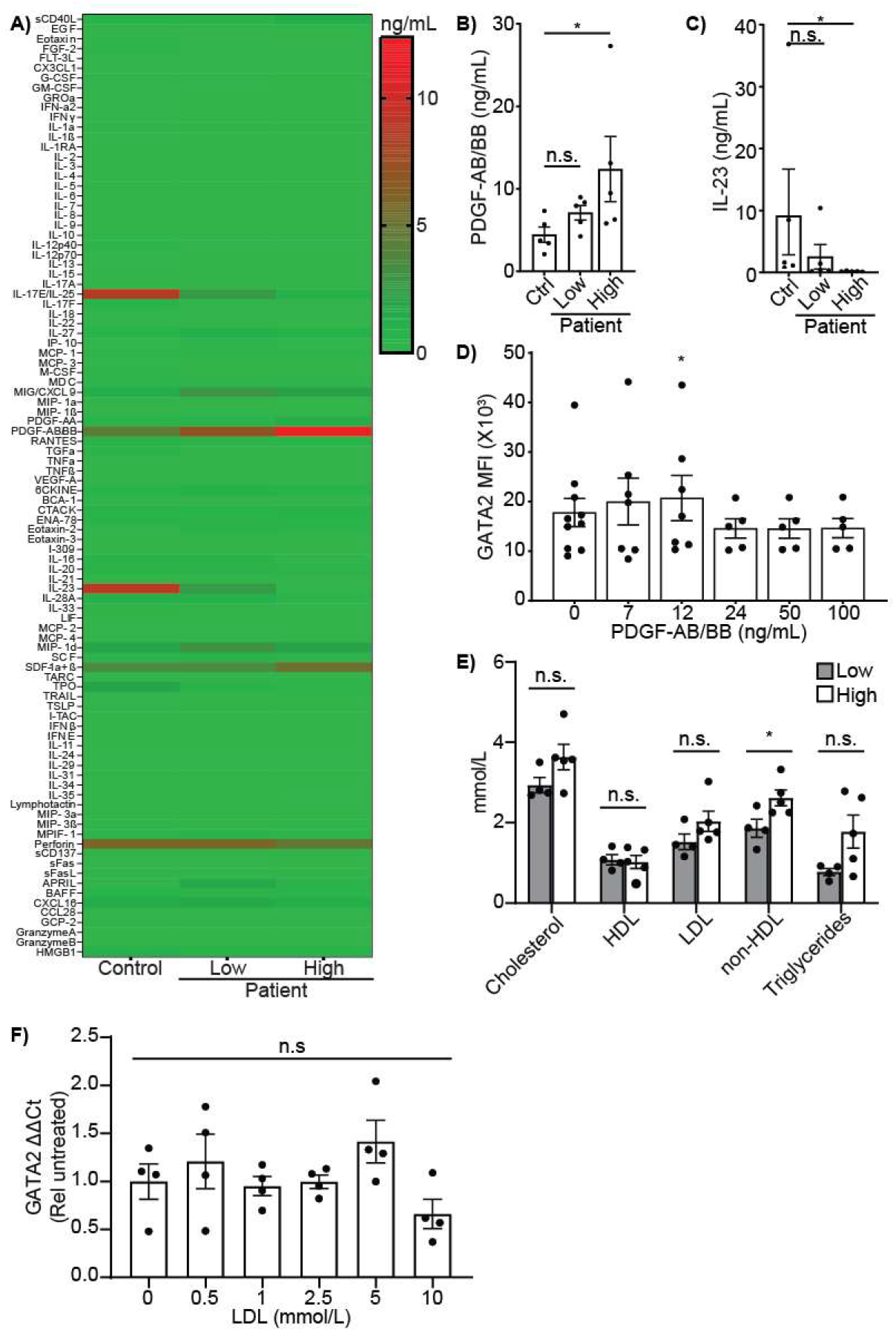
Plasma cytokines and lipids do not drive GATA2 expression. A 96-plex Luminex panel was used to quantify cytokine concentrations in the serum of healthy controls versus atherosclerosis patients with high (MFI >2 S.D. compared to healthy controls, “High”) or low monocytic GATA2 expression. **A)** Heatmap of serum concentrations of all 96 cytokines in healthy controls versus patients with low or high monocytic GATA2 expression. **B-C)** Serum concentration of PDGF-AB/BB (B) and IL-23 (C), the only two cytokines significantly different in GATA2-High patients versus healthy controls. **D)** Expression of GATA2 in monocyte-derived macrophages (moMacs) differentiated in the presence of the indicated concentration of PDGF-AB/BB for 48 hours. **E)** Serum concentrations of total cholesterol (Cholesterol), HDL, LDL, non-HDL cholesterol, and total triglycerides (Triglycerides) in atherosclerosis patients with high or low monocytic GATA2 expression. **F)** Expression of GATA2 in moMacs stimulated with the indicated concentrations of LDL for 3 days. ΔΔCt values are normalized to the mean ΔΔCt of untreated moMacs (0 mmol/L LDL). n = 3 (A) or n ≥ 4 (B-F), * = p < 0.05 between indicated groups (B,C,E) or 0 ng/mL PDGF-AB/BB (D), n.s. = p > 0.05, Kruskal-Wallis test with Dunn correction (B-C, F), Dunnett’s test (D), or Mann-Whitney U test (E).

### GATA2 Drives Macrophage Proliferation Throughout Disease

Up to this point, all analysis of GATA2 expression in plaque macrophages was performed in early-stage atheromas present in aortic biopsies acquired during coronary artery bypass graft surgery. As these are early-stage or arrested plaques (Stary scores of 1 to 2, [34]), they may not represent GATA2 expression patterns throughout plaque development. Moreover, the gene expression profile of these cells suggests that they may be proliferating. As such, we used histological staining and fluorescence immunohistochemistry of human coronary vessels with advanced plaques to identify macrophages (CD68+ cells), GATA2, and the cell proliferation marker Ki-67 (**Figure 4A**). Using a trained machine learning algorithm [35], we identified the macrophages in each lesion and classified them based on their expression of GATA2 and Ki-67, finding that the GATA2+/Ki-67+ proliferating population was enriched near the lesion borders (**Figure 4B**, yellow cells). There was no correlation between lesion size and the portion of lesional cells that were macrophages, but there was a modest correlation between the density of macrophages in the lesion and the lesion size (**Figure 4C, D**). Ten percent of cells in the atheroma were proliferating, with macrophages accounting for 15% of the proliferating population (**Figure 4E**), with between 92% to 100% of proliferating macrophages expressing GATA2 (**Figure 4F, G**). Interestingly, most of the proliferating non-macrophage cells were also GATA2+, indicating that GATA2 upregulation is correlated with the proliferation of multiple cell types in atherosclerotic lesions (**Figure 4B**, tan cells). These data indicate that GATA2 expression is concomitant with plaque macrophage proliferation, although it is unclear if GATA2 is driving proliferation, or if GATA2 expression is a product of proliferation.

**Figure 4:**
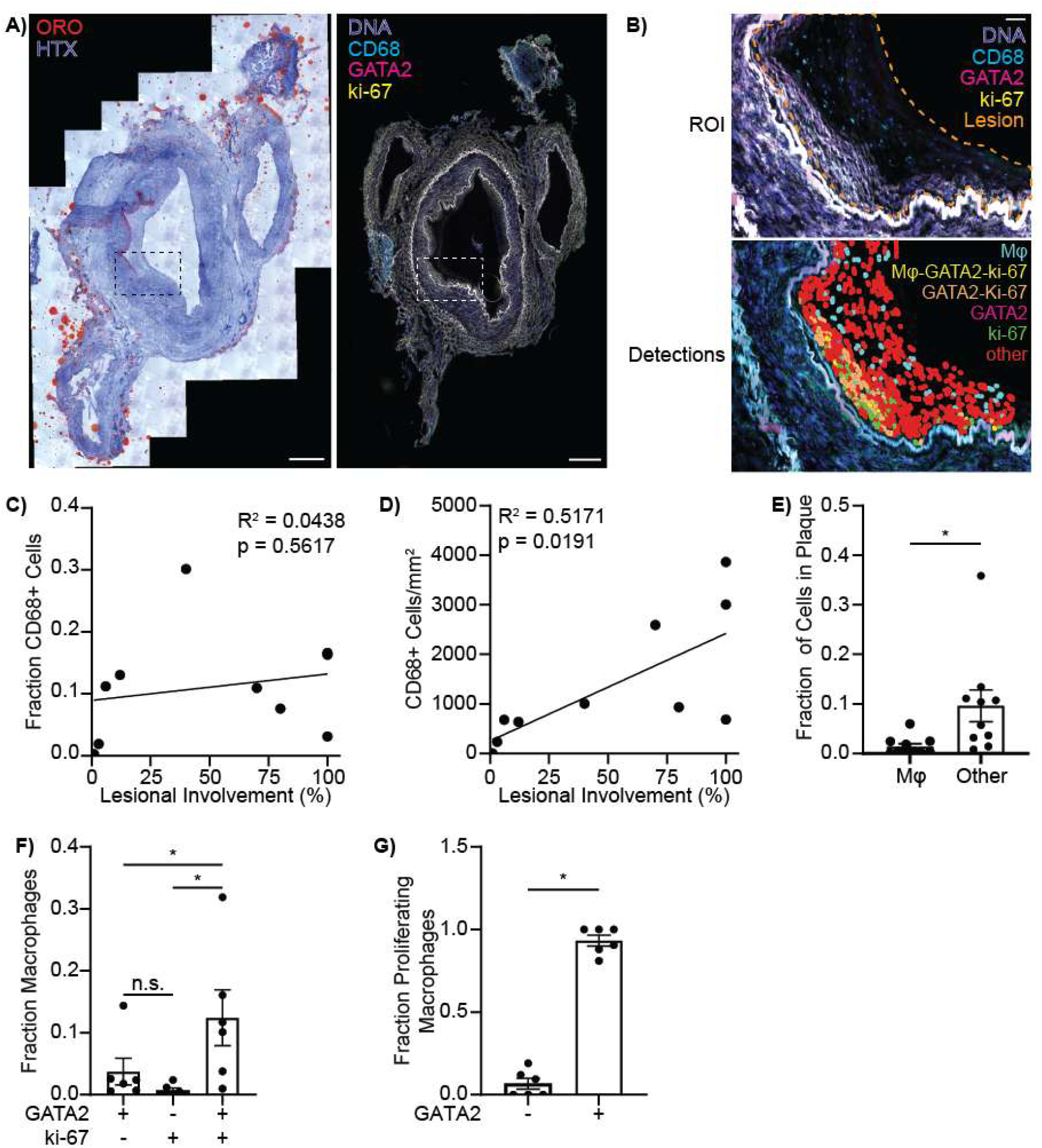
GATA2 is Correlated with Macrophage Proliferation in Advanced Atherosclerotic Plaque. Fluorescence histology of human coronary blood vessels was used to identify proliferating cells using the marker Ki-67, macrophages with the marker CD68+, and GATA2. **A)** Serial sections of the same coronary blood vessel showing lipids (Oil-Red-O/ORO) and cellularity (Hematoxylin/HTX, *left*), and fluorescent labeling (*right*) for DNA (blue), macrophages/CD68 (cyan), GATA2 (magenta) and Ki-67 (yellow). Scale bars are 500 μm, boxed area indicates a lesional area shown in higher magnification in panel B. **B)** Magnified area of the region of interest (ROI) from panel A. Top: fluorescent labeling for DNA (blue), macrophages/CD68 (cyan), GATA2 (magenta) and Ki-67 (yellow). The lesional area is indicated by the orange dotted line. Bottom: Automated classification of all cells in the lesional area. Cells are classified based on their expression of CD68 (macrophages), Ki-67, and GATA2. Double-negative macrophages (Mφ) are in cyan, GATA2+/ki67+ macrophages (Mφ-GATA2-Ki-67) are in yellow; no single-positive macrophages are present. Non-macrophages (CD68-cells) expressing GATA2 (magenta), Ki-67 (green), both GATA2 and Ki-67 (tan), or neither marker (red) are also indicated. Scale bar is 50 μm. **C)** There is no correlation between the portion of the blood vessel containing atherosclerotic plaque (Lesional Involvement) and the fraction of cells in the lesions that are macrophages (CD68+ cells). **D)** The density of macrophages in a plaque is positively correlated with the portion of the blood vessel containing atherosclerotic plaque (Lesional Involvement). **E)** Quantification of the portion of dividing cells (Ki-67+ cells) within lesions that are macrophages (Mφ, CD68+ cells) versus other cell types (CD68-cell types). **F)** Quantification of the fraction of macrophages in atherosclerotic lesions that express GATA2, Ki-67, or both markers. **G)** Portion of proliferating macrophages which lack (−) or express (+) GATA2. n = 9, * = p < 0.05, n.s. = p > 0.05 between indicated groups, Mann-Whitney U test (E,G) or Kruskal-Wallis test with Dunn correction (F). Correlative R2 and p values were calculated using a Spearman’s correlation coefficient. Fluorescent images in panels A and B have had their intensity square root-normalized to better illustrate the distribution of cells within the images.

### GATA2 IS Required for Macrophage Proliferation

To determine if GATA2 was required for macrophage proliferation, we first performed a network interaction analysis of genes expressed in GATA2+ macrophages and identified a potential proliferation-regulating network that was enriched in these cells (**Figure 5A**). This network was reminiscent of the same proliferative network mediated by GATA2 during myelopoiesis, particularly in the upregulation of the protooncogene MYB, and in the downregulation of JUN—necessary for avoiding aberrant monocyte differentiation [32,36]. We utilized GATA2 over-expressing human THP1 monocytes to investigate the impact of GATA2 expression on this putative pathway, allowing us to investigate GATA2’s role in the absence of additional signaling induced by oxLDL or other plaque-derived signals. Consistent with the network analysis, GATA2 overexpression upregulated MYB and reduced expression of JUN (**Figure 5B**). We next used a luciferase reporter assay to determine if GATA2 was directly regulating expression of JUN and MYB during macrophage differentiation. Unexpectedly, MYB promoter activity did not increase when GATA2 was overexpressed, whereas JUN promotor activity was strongly suppressed (**Figure 5C-D**), indicating that GATA2 directly regulates the JUN promoter, but may be regulating MYB expression via distal or intronic regulatory elements not included in our luciferase assay [32,37,38]. Using a dye-dilution assay, the proliferation of THP1 monocytes during macrophage differentiation was quantified, with GATA2 overexpressing cells proliferating significantly more than wild-type cells (**Figure 5E, S3)**. These results indicate that the expression of GATA2 is sufficient to drive macrophage proliferation, but it is unclear whether this occurs under atherogenic conditions. GATA2 was upregulated in oxLDL-stimulated THP1 macrophages (**Figure 5F**), with oxLDL stimulation inducing proliferation of THP1 cells during macrophage differentiation (**Figure 5G**). Critically, this proliferation was suppressed in THP1 cells expressing a GATA2-targeting shRNA, or when THP1 cells were treated with a cell-penetrant siRNA against MYB (**Figure 5G, S3**). Lipid-laden macrophages often undergo apoptosis due to activation of the unfolded protein response triggered by the intracellular accumulation of sterols and lipids. Our gene expression analysis of the GATA2+ macrophage transcriptome identified upregulation of several anti-apoptotic genes (**Figure 1G)**, suggesting that GATA2 expression may desensitize macrophages to apoptosis. While we did not observe apoptosis in response to oxLDL, GATA2 overexpression reduced apoptosis, and GATA2 knockdown increased apoptosis, following induction of the unfolded protein response with thapsigargin (**Figure 5H)** [39]. Combined, these results indicate that GATA2 is expressed in response to oxLDL, thereby promoting atheroma development by inducing macrophage proliferation and desensitizing these cells to apoptotic stressors present in the atheroma.

**Figure 5:**
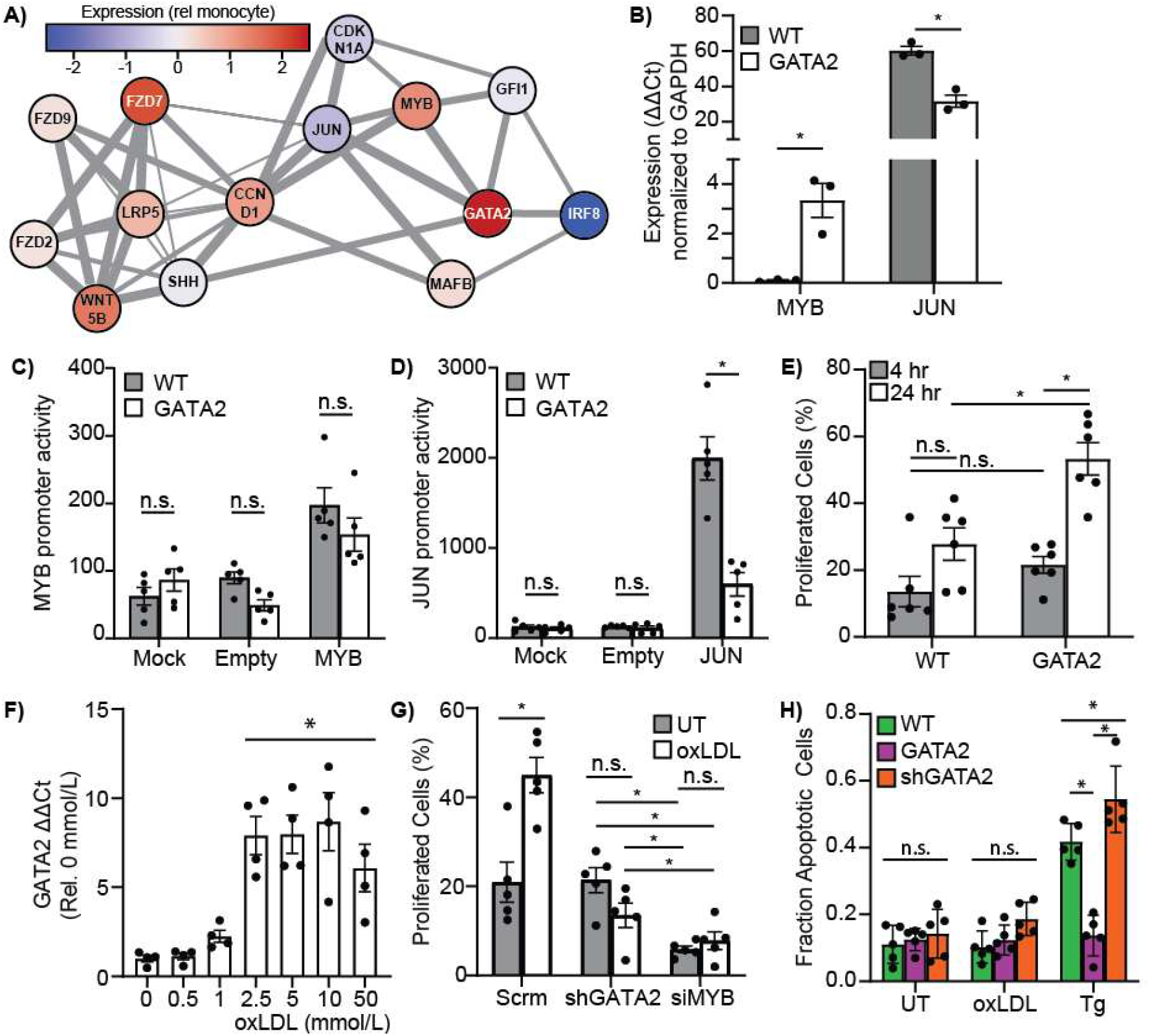
GATA2 Drives Macrophage Proliferation. **A)** Cytoscape network analysis identified enrichment of a proliferative network in the transcriptome of GATA2+ macrophages compared to monocytes from healthy controls. **B)** Overexpression of GATA2 in THP1-derived macrophages increases the expression of MYB and decreases the expression of JUN compared to wild-type THP1 macrophages. **C-D)** Quantification of MYB (C) and JUN (D) promoter activity using firefly luciferase reporters containing the MYB or JUN promoters, in wild-type (WT) versus GATA2-overexpressing (GATA2) THP1-derived macrophages. Cells are either untransfected (Mock), transfected with a promoterless vector (Empty) or transfected with a MYB- or JUN-promoter containing vector. **E)** Impact of GATA2 on the proliferation rate of GATA2-overexpressing (GATA2) versus wild-type (WT) THP1 monocytes as they differentiate into macrophages. **F)** GATA2 expression in THP1-macrophages following exposure to the indicated concentrations of oxLDL for 48 hours. **G)** Impact of GATA2 and MYB siRNA knockdown, compared to a scrambled siRNA control (siRNA), on the oxLDL-induced proliferation of THP1-macrophages. **H)** THP1-derived macrophage apoptosis under control conditions (UT) versus following stimulation with 2.5 mmol/L oxLDL (oxLDL), or 10 μM thapsigargin (Tg). Macrophages are either derived from wild-type THP1 cells (WT, green), GATA2-overexpressing THP1 cells (GATA2, magenta), or THP1 cells expressing a GATA2-depleating shRNA (shGATA2, orange). n = 4 – 6, * = p < 0.05, n.s. = p > 0.05, between indicated groups, Mann-Whitney U test (B-E, G) or Kruskal-Wallis test with Dunn correction (F, H).

## Discussion

Macrophage proliferation produces the majority of macrophages present in an atherosclerotic plaque, suggesting that reducing macrophage proliferation could be a potentially powerful therapeutic approach for treatment of atherosclerosis [9,10]. Using human aortic biopsies and coronary artery sections, we identified the hematopoietic transcription factor GATA2 as a protein uniquely expressed in proliferating plaque macrophages. These GATA2+ macrophages had a transcriptional profile with characteristics of both monocytes and macrophages, indicating that these cells were either arrested at a mid-point in monocyte differentiation, or were macrophages which had partially de-differentiated to an immature state. These cells expressed the proliferation marker Ki-67, and *in vitro*, GATA2 expression was necessary and sufficient to induce macrophage proliferation, mediating oxLDL-induced proliferation by upregulation of the protooncogene MYB. Together, these results lead us to conclude that oxLDL-induced GATA2 expression in atherosclerotic plaque macrophages induces macrophage proliferation via a GATA2-MYB mediated proliferative network. Thus, GATA2 or its downstream effectors may be viable therapeutic targets for the treatment of atherosclerosis, given that the bulk of macrophages in the plaque arise via proliferation [9].

Previous studies have linked single nucleotide polymorphisms (SNPs) in GATA2 to atherosclerosis, although there does not appear to be a consistent mechanism in how different SNPs contribute to disease. Muiya *et al*. identified 8 SNPs in GATA2 linked to coronary artery disease, with broadly different impacts on disease risk factors and disease progression. This included identification of SNPs associated with LDL levels, type 2 diabetes, obesity, hypercholesterolemia, and predisposition to myocardial infarct without impacts on other prognostic markers [40]. Only one of these SNPs affected the coding region of GATA2 (rs2335052), with the remainder located in the intron between exons 5 and 6, in the 3’ UTR, or in the non-transcribed region 3’ to the GATA2 gene. This coding sequence SNP is associated with an increased risk of coronary artery disease and some cancers, and results in an Ala^164^Thr mutation in the negative regulatory domain of GATA2, resulting in increased GATA2 protein levels [41– 44]. Muiya *et al*. also identified a common frameshift mutation that results in the haploid expression of GATA2, but no link to atherosclerosis was found even though the same mutation predisposes people to leukemia [40,45]. Other mutations that decrease GATA2 expression, such as those causing MonoMac syndrome, are also not associated with atherosclerosis [46]. GATA2 activity and expression levels can also be controlled via post-translational modifications. For example, GATA2 phosphorylation reduces the nuclear localization of the GATA2 protein, with adipocytes expressing a form of GATA2 that cannot be phosphorylated (e.g. constitutively nuclear localized) producing significantly elevated levels of pro-atherogenic cytokines including MCP-1 and GM-CSF [47–49]. While the mechanisms by which these various SNPs and pathways drive atherosclerosis are unclear, these studies consistently show that transcriptional and post-translational modifications that increase GATA2 expression or activity are atherogenic, while no linkage is observed between atherosclerosis and processes that reduce GATA2 expression or activity. These findings are consistent with our observations, wherein elevated GATA2 expression drove pro-atherogenic changes in macrophage proliferation and sensitivity to apoptosis.

In adults, GATA2 expression is generally limited to two sites: lymphatic endothelium and to hematopoietic stem cells and myeloid progenitors in the bone marrow [50,51]. In both locations, GATA2 is critical for the self-renewal of these cell types, with GATA2 expression driven by cyclin-dependent kinases [31]. GATA2 expression is self-reenforcing via a positive feedback loop in which GATA2 activates its own promoter [52–54]. In the bone marrow, GATA2 expression must be reduced for myeloid cells to enter terminal differentiation. This cessation of GATA2 expression is mediated by a GATA2/GATA1 switch, wherein upregulation of GATA1 displaces GATA2 from its binding sites in the GATA2 promoter, leading to decreased GATA2 expression [54,55]. Other than our initial identification of GATA2 in atherosclerotic plaque macrophages, there are no reports of human GATA2 expression in non-cancerous peripheral myeloid cells [18]. Indeed, the only report of GATA2 expression in mature macrophages is in the alveolar macrophages of rats chronically infected with the fungal pathogen *Pneumocystis carinii*, where GATA2 expression impairs macrophage phagocytosis [56]. In mouse macrophages, toll-like receptor 4 signaling, triggered by LPS, promotes GATA2 expression which then upregulates expression of the inflammatory cytokine IL-1β, with MYD88 and MEK1/2 signaling initiating the nuclear translocation of GATA2 [57]. These results suggest that GATA2 expression in peripheral macrophages is an aberrant state, perhaps brought on by persistent inflammation or other non-resolving immunological stimuli. While not detected in myeloid cells, human lung endothelial cells upregulate GATA2 in response to acute infection with SARS-CoV-2 where it contributes to the production of inflammatory cytokines, again consistent with GATA2 expression being induced by inflammation [58,59]. GATA2 expression is also induced by cell stress, with increased activity of the GATA2 promoter observed after heat stress, and GATA2 expression promoting reendothelialization of injured carotid arteries [38,60]. Our observation that oxLDL, but not LDL, potently induced GATA2 expression in macrophages is consistent with both of these processes, as oxLDL induces inflammatory signaling via CD36 and toll like receptors 2 and 4, as well as provokes cell stress via its intracellular accumulation and activation of the unfolded protein response [4,61].

We found that oxLDL-induced proliferation required both GATA2 and MYB, which is consistent with previous studies that identified MYB as a transcriptional target of GATA2, and which demonstrated that MYB was required for GATA2-mediated transition of cells from G_1_ to S phase [32]. While we saw a significant induction of MYB mRNA in cells overexpressing GATA2, we did not see a concurrent increase in MYB promoter activity. This was not unexpected as GATA2 and other GATA transcription factors regulate MYB largely via distal promoter elements and via GATA2 binding to intronic sites located within the MYB gene – regions not captured in our luciferase vectors [32,62]. We also observed downregulation of JUN promoter activity and mRNA levels. While JUN can mediate progression through G_1_ phase, via regulating cyclin D1 levels, JUN is antagonistic to GATA2 activity [36,63]. While little research has been done into the regulation of JUN by GATA-family transcription factors, ChIP-seq analyses have identified a site bound by GATA2, GATA3, GATA4 and GATA6 in the JUN promoter, consisting of a canonical AGATAA binding motif, located roughly 60 base-pairs 5’ to JUN’s transcriptional start site [64]. While the mechanism by which GATA2 suppresses JUN expression is not clear from our data, GATA transcription factors are known to engage in “switch” behaviour, where GATAs that bind to the same site displace each other in a competitive manner that is dependent on the relative affinity for the binding site and the relative abundance of the GATA proteins [65–67]. Thus, increased expression of GATA2 could displace a normally activating GATA at this site, with GATA2 then negatively regulating the expression of JUN. Given that GATA2 expression is lowest at the start of G_1_ phase, and increases as cells enter the G_1_/S transition and pass through to G_2_ phase, these data indicate that GATA2 and JUN are upregulated at different points of the cell cycle, with cross-talk between these genes controlling both their mutual expression, and controlling cell cycle progression [31].

In summary, we have identified GATA2 as a transcription factor upregulated in a subpopulation of macrophages present throughout atheroma development. These GATA2+ macrophages feature a transcriptome mid-way between monocytes and fully differentiated macrophages, and are the dominant population of replicating macrophages in the plaque. In these cells, GATA2 induces cell cycle progression via upregulation of the protooncogene MYB, while simultaneously decreasing these cells sensitivity to the induction of apoptosis induced by cell stress pathways. Given that most macrophages present in atherosclerotic plaque arise via macrophage proliferation, this GATA2-mediated pathway likely accounts for much of the cellularity of plaques and represents a potential therapeutic target for reducing plaque burden and inducing plaque regression.

## Materials & Methods

### Materials

pGL4.20[luc2/Puro] was a gift from Dr. Rodney DeKoter, University of Western Ontario. Lympholyte poly and THP1 human monocytic cells (ATCC TIB 202) were purchased from Cedarlane Labs (Burlington, Canada). THP1 cells transduced with a GATA2 overexpression cassette, or GATA2-targeting shRNA, were generated previously [18]. Information on all antibodies used in this study can be found in **Supplemental Table 1**. Information on all primers used in this study can be found in **Supplemental Table 2**; all primers and synthetic DNA blocks were from IDT (Coralville IA). Human TruStain Fc Block and the CD14 cell Mojosort enrichment kit were from Biolegend (San Diego, CA). FlowJo software, blood collection sets, and heparinized vacuum tubes were purchased from Becton-Dickinson (Franklin Lakes, NJ). Aortic punches were purchased from QUEST Medical (London, Canada). OCT freezing compound and tissue cassettes were from Sakura Finetek (Torrance, CA). DMEM, RPMI, 100× antibiotic/antimycotic, and fetal bovine serum (FBS) were purchased from Wisent (Saint-Jean-Baptiste, Canada). Recombinant human M-CSF, PDGF-AA/AB, TNFα, and IL-23 were from Peprotech (Rocky Hill, NJ). Cholesterol and triglyceride quantification kits and Oil-Red-O were from Abcam (Cambridge, UK). 96-well luciferase plates were from Greiner Bio-One (Kremsmünster, Austria). Coverslips, 16% paraformaldehyde (PFA), and immersion oil were from Electron Microscopy Sciences (Hatfield, PA). Human Coronary Artery Endothelial Cells and endothelial cell culture media were from Promocell (Heidelberg, Germany). Ibidi μ-Slide VI^0.4^ flow chambers were from Ibidi GbmH (Gräfelfing, Germany). HiFI Gibson assembly mix, restriction enzymes, T4 DNA ligase, and Q4 PCR mastermix were purchased from NEB Canada (Whitby, Canada). Amaxa nucleofector kits were purchased from Lonza Canada (Kingston, Canada). Luciferase detection kits were from Promega (Madison WI). Accell cell-penetrant siRNAs and Accell medium was from Horizon Discovery (Waterbeach, UK). RNA isolation kits were from Qiagen (Hilden, Germany). iScript cDNA kits and SsoFast Evagreen Supermix were from Biorad (Hercules, CA). Prism software was from Graphpad Software (Boston, MA). Common laboratory chemicals were purchased from Bioshop Canada (Mississauga, Canada). All other materials were purchased from ThermoFisher (Waltham, MA).

### Ethics & Sample Collection

This study used blood and tissue samples from patients undergoing coronary artery bypass graft surgery at University Hospital, London, Ontario, Canada. The study was reviewed and approved by the Human Research Ethics Committee of the Western University Health Sciences Research Ethics Board (HSREB approval numbers: 107566 and 104010). Although discarded tissue was used in this study, consent was obtained from patients before enrolling. Participants’ age range was from 48 to 78 years, and they were 18% female and 82% male. Blood was collected from patients into heparinized tubes during pre-operative procedures. Aortic punch tissue was obtained intra-operatively using a 4.0 mm diameter CleanCut RCL Aortic Punch and immediately placed into cold saline. This discarded aorta tissue was then halved, with one half stored at -80°C after flash freezing in liquid nitrogen, and the other embedded in OCT freezing compound for sectioning. Blood was collected from healthy age- and sex-matched controls by venipuncture and collected into heparinized tubes.

### Peripheral Blood Monocyte and Plasma Isolation

Peripheral blood mononuclear cells were isolated from blood using density gradient centrifugation. Initially, blood was centrifuged at 400 × g for 10 min to separate plasma. Plasma was collected from the top of the tube, and the remaining pellet diluted with a volume of PBS equal to the starting volume of blood. 3-4 mL of lympholyte poly was placed into a 15 mL tube, and the diluted blood layered on top. This tube was centrifuged at 500 × g for 25 min with medium acceleration and no brake. The top band of mononuclear cells was carefully collected into a 50 mL tube and washed with 50 mL phosphate buffered saline (PBS, 137 mM NaCl, 10 mM Na_2_HPO_4_, pH 7.4), then centrifuged at 400 × g for 5 min with maximum acceleration and braking. After washing, cells were suspended in 10 mL PBS and counted. In some experiments, monocytes were purified from this mixed population by positive-selection magnetic sorting using a Mojosort kit as per the manufacture’s instructions. Cells were then washed again at 400 × g for 5 min and suspended in either culture medium (RPMI + 10% FBS + 1:100 antimycotic/antibiotic) for immediate use, or were frozen in freezing medium (culture medium + 10% dimethyl sulfoxide) for later experiments.

### Differentiation of Monocyte-Derived Macrophages

Macrophages were differentiated from peripheral blood mononuclear cells using either cytokines or autologous plasma. 2.5 × 10^5^ peripheral blood mononuclear cells in culture medium were placed into each well of a 12-well tissue culture plate and incubated in a 5% CO_2_/37°C incubator for 1 hr, then non-adherent cells removed with two gentle washes with pre-warmed PBS. For cytokine differentiation, the final PBS wash was replaced with culture medium + 10 ng/mL human M-CSF. The cells were incubated for 3 days in a 5% CO_2_/37°C incubator, then the medium replaced with fresh culture medium + 10 ng/mL human M-CSF, and three days later the cells used for experiments. For autologous plasma differentiation, the final PBS wash was replaced with culture medium +10% autologous plasma. The cells were incubated for 3 days in a 5% CO_2_/37°C incubator, then the medium replaced with fresh culture medium + 10% autologous plasma, and three days later the cells used for experiments.

### Transcriptomics Analysis

Transcriptomes of GATA2+ macrophages, purified by laser microdissection from aortic punch biopsies, and moMacs derived from age- and sex-matched controls were published previously (Borealis dataset WYSAA0, https://doi.org/10.5683/SP2/WYSAA0) [18]. Healthy donor monocyte transcriptomes were acquired from Farnia *et al*., GEO accession GSE86984 [68]. Analysis was performed in Bioconductor [69]. First, raw microarray data was imported, converted to expression values, and quality confirmed via the arrayQualityMetrics command. Robust multichip average was then calculated for all transcripts, and differentially expressed genes identified as those with >2-fold change in expression between any two groups. Principal component analysis was performed on the differentially expressed genes, and a gene expression heatmap prepared by calculating the Manhattan distance between all differentially expressed genes that was then plotted using the pheatmap command. The genes from individual gene clusters were extracted, and their enrichment into reactomes calculated using ReactomePA. Network analysis for transcriptional networks was performed using Cytoscape [70].

### Flow Cytometry

Sorted CD14+ cells or whole PBMCs were first blocked with Human TruStain FcX blocker for 10 min at room temperature. After washing, cells were stained with human anti-CD14-APC antibody and fixable viability dye-eFluor780 for 45 min at 4°C. For intranuclear GATA2 and Ki-67 staining, the cells were fixed and permeabilized with flow cytometry permeabilization solution (ThermoFisher), followed by addition of human anti-GATA2-AF555 and anti-Ki-67-FITC antibodies for 1 hr at room temperature. After washing, the cells were resuspended in FACS buffer (2% FBS in PBS) and a minimum of 10,000 cells acquired on a BD FACSCanto II. FlowJo was used to analyze the resulting data by first gating on forward and side scatter, selecting singlets with the FSC-H/-A channels, live cells gated via the eF780 channel, and monocytes identified as CD14+ cells in the gated population. The mean fluorescence intensity for GATA2 and Ki-67 was then determined for the CD14+ cell population. The specific antibodies and staining conditions used for flow cytometry are listed in **Supplemental Table 1**.

### Luminex Cytokine Assays

Plasma samples were platelet-depleted by centrifuging at 21,000 × g for 30 sec. 100 μL of the clarified plasma was then frozen and sent to Eve Technologies Corporation (Calgary, Canada) for analysis on a Luminex 200 system and Eve Technologies’ Human Cytokine 96-Plex Discovery Assay. Quantified cytokines included: sCD40L, EGF, Eotaxin, FGF-2, FLT-3 Ligand, Fractalkine, G-CSF, GM-CSF, GROα, IFN-α2, IFN-γ, IL-1α, IL-1β, IL-1RA, IL-2, IL-3, IL-4, IL-5, IL-6, IL-7, IL-8, IL-9, IL-10, IL-12(p40), IL-12(p70), IL-13, IL-15, IL-17A, IL-17E/IL-25, IL-17F, IL-18, IL-22, IL-27, IP-10, MCP-1, MCP-3, M-CSF, MDC, MIG/CXCL9, MIP-1α, MIP-1β, PDGF-AA, PDGF-AB/BB, RANTES, TGFα, TNF-α, TNF-β, VEGF-A, 6CKine, APRIL, BAFF, BCA-1, CCL28, CTACK, CXCL16, ENA-78, Eotaxin-2, Eotaxin-3, GCP-2, Granzyme A, Granzyme B, HMGB1, I-309, I-TAC, IFNβ, IFNω, IL-11, IL-16, IL-20, IL-21, IL-23, IL-24, IL-28A, IL-29, IL-31, IL-33, IL-34, IL-35, LIF, Lymphotactin, MCP-2, MCP-4, MIP-1δ, MIP-3α, MIP-3β, MPIF-1, Perforin, sCD137, SCF, SDF-1, sFAS, sFASL, TARC, TPO, TRAIL, and TSLP.

### Cholesterol and Triglyceride Quantification

Total cholesterol, HDL, and LDL cholesterol were measured in the patient’s plasma using a cholesterol assay kit. For total cholesterol, whole plasma was used, whereas HDL and LDL were quantified using a 1:1 mixture of plasma and 2× Precipitation Buffer. Plasma was centrifuged for 10 minutes at 2,000 × g and the supernatant used for HDL-cholesterol measurement, while the precipitate was resuspended in 100 μL PBS for measuring the LDL/VLDL fraction. 50 μL volumes of reaction mix were added to the total cholesterol, HDL-cholesterol, LDL-cholesterol, and standard curve samples according to the manufacturer protocol. The mixture was incubated for 60 min at 37°C and read on a microplate reader with excitation at 535 nm and emission at 587 nm. Triglycerides were measured in patient’s plasma using a triglyceride assay kit. 2 μL of cholesterol esterase/lipase was added to 10 μL aliquots of plasma and incubated for 20 min at RT with constant agitation. Next, triglyceride assay buffer, OxiRed probe, and triglyceride enzyme mix were added to a final volume of 50 μL in 96-well plates and incubated for 60 min at RT. The plate was read on a microplate reader with an excitation of 535 nm and an emission of 587 nm.

### Parallel Plate Flow Chamber

A parallel plate flow chamber was used to simulate aortic monocyte recruitment on primary human aortic endothelial cells. Human Coronary Artery Endothelial Cells from an 84-year-old male patient were seeded in endothelial cell growth medium into an Ibidi μ-Slide VI^0.4^ flow chamber. Cells were cultured at 37°C/5% CO_2_ until confluent, typically 48-72 hours. To mimic plaque inflammation, the media was replaced with medium containing 5 ng/mL human TNFα, followed by incubation for 4 hr at 37°C/5% CO_2_. The μ-Slide VI^0.4^ was placed on the heated stage of a Leica DMI600B microscope equipped with an Inscoper control unit (Inscoper, Cesson-Sévigné, France), and imaged with DIC illumination using a 40×/1.30 NA oil immersion lens. During imaging, sorted CD14+ cells were perfused through the flow chamber with an Orion Sage M362 syringe pump at a rate of 39.5 mL/min, producing a surface shear force of 50 dynes/cm^2^. Timelapses of monocyte recruitment were captured for 30 min, imaging every 30 sec. The resulting images were analyzed in FIJI to quantify monocyte rolling, adhesion, and transmigration across the endothelium as described previously [71,72].

### Tissue Imaging

For aortic biopsies, OCT embedded tissues were sliced into 8 μm thick sections using a Leica CM18602 cryostat, and slices placed onto glass slides. For advanced atheromas, pre-sliced 10 μm thick sections of pathologist-scored human coronary arteries from atherosclerosis patients were purchased from Origene, selecting patients with a range of disease severity. Slides were kept frozen at -80°C until processed. For Oil-Red-O staining, the tissue was fixed in ice-cold 10% formalin for 10 min, rinsed in 3 changes of distilled water, air dried, then immersed in propylene glycol for 5 min. The tissue was then stained in pre-warmed 0.5% Oil Red O solution in propylene glycol for 10 min in a 60oC oven. Following staining, the tissue was destained in 85% propylene glycol in water for 5 min and rinsed with 2 changes of distilled water. For counter-staining, the tissue was treated with Mayer’s hematoxylin for 30 sec, washed under running water for 3 min, and placed in distilled water for an additional 3 min. The stained tissue was then mounted on a coverslip using an aqueous mounting medium and imaged. For fluorescence staining, samples were thawed to RT, fixed in fresh 4% PFA in PIPES buffer (0.1M PIPES, 5 mM EGTA, 2mM MgCl2 · 6H_2_O) for 10 min at 37°C, then washed with PBS three times in Coplin Jars. Samples were blocked with PBS + 10% Normal Donkey Serum + 0.1% Triton X-100 + 0.01% NaN_3_ at room temperature for 1 hr. Samples were stained with primary anti-CD68 and anti-Ki67 antibodies for 1.5 hours at 37°C. After three washes in fresh PBS, the samples were stained with 0.67 ug/ml of anti-rabbit 488 and anti-mouse 647 secondary antibodies for 45 minutes at RT. The samples were washed three times with PBS and stained with anti-GATA2-AF555 for one hour at RT. To stain nuclei, 1 μg/mL of Hoechst in PBS was added and incubated at room temperature for 10 min. Following a final PBS wash, a coverslip was mounted on top of the samples with an aqueous mounting medium. Samples were imaged using a 20×/0.75 NA objective on a Leica DMI 5500B microscope equipped with a halogen light source, filters for DAPI, FITC, Cy3 and Cy5 fluorophores, operated using LAS-AS software. Histological samples were imaged using a colour camera and brightfield illumination, whereas fluorescence samples were imaged using the appropriate filter sets. Fluorescent samples were then 2D deconvolved using a 30-iteration Richardson-Lucy algorithm in DeconvolutionLab2 [73]. To perform a quantitative analysis of macrophage phenotypes, blinded images were used to train the QuPath ML algorithm [35], first training the algorithm to segment nuclei, macrophages (CD68+ cells), followed by training to classify cells based on their expression patterns of Ki-67 and GATA2. The trained algorithm was then used to quantify all tissue sections. Detections and classifications were then exported for graphing and statistical analysis.

### THP1 Cell Culture

Wild-type THP1 human monocytes, and THP1 cells ectopically over-expressing GATA2, or expressing a GATA2-targeting shRNA, were maintained as suspension cultures in RPMI + 10% FBS in a 37°C/5% CO_2_ incubator. Cells were grown to a density of 1 × 106/mL and then split by diluting 1:5 into fresh medium. To differentiate the cells into macrophages, 2.5 × 105 cells were placed into each well of a 12-well plate, and the cells incubated in RPMI + 10% FBS + 100 ng/mL PMA for 72 hr. For experiments involving microscopy, #1.5 thickness, 18 mm diameter circular coverslips were cleaned by acid washing (2 M HCl in ddH_2_O, 56°C, overnight), autoclaved, and then placed into the wells of the 12-well plate prior to plating cells.

### Luciferase Assays

The MYB and JUN promoters were ordered as synthetic DNA constructs. The promoters were inserted into pGL4.20[luc2/Puro] vector via Gibson assembly by first linearizing the vector with an EcoRV digest, followed by gel purification, and assembly of the final vector using a HiFi Gibson Assembly Kit. Successful cloning was confirmed by Sanger sequencing at the London Genomics Centre (London, Canada). Wild type THP-1 or GATA2-overexpressed THP-1 cells were transfected with 0.5 μg of DNA using an Amaxa Human Monocyte Nucleofector kit and a Lonza Nucleofector II, using the manufacturer’s V-001 protocol. After transfection, 500 μL of warm RPMI + 10% FBS was added to the electroporation cuvette. Transfected cells were then transferred to a 12-well plate and incubated in a humidified 37^°^C/5% CO_2_ incubator overnight. The next day, macrophage differentiation was induced, as described above. Following differentiation, the cells were lysed with 250 μL passive lysis buffer by rocking for 20 mins at room temperature. Lysates were collected centrifuged at 21,000 × g for 30 sec to remove debris, and 100 μL of lysate added to each well of a 96-well white polystyrene plate. 50 μL of luciferase assay reagent II was added to the well, and the plate immediately read using 2(L) Lum, linear shaking, using a 10 sec acquisition in Cytation Gen5 software.

### Proliferation Assays

To quantify proliferation during macrophage differentiation, THP-1 monocytes were pre-stained with Cell Proliferation Dye eFluor 670 by suspending the cells in PBS + 5 μM Cell Proliferation eFluor 670 dye for 5 min, followed by pelleting the cells with a 400 × g centrifugation and resuspending the cells in RPMI + 10% FBS + 100 ng/mL PMA. For quantification of macrophage proliferation, THP1 macrophages were differentiated as above, treated for 24 hr with 1 μM of either a non-targeting (scrambled) or MYB targeting Accell cell-penetrant siRNAs in Accell medium, cultured for an additional 24 hr in RPMI + 10% FBS, then stimulated with 2.5 mmol/L (∼100 μg/mL) oxLDL. Immediately after PMA or oxLDL stimulation an 11×11 tiled region was imaged in DIC and far-red fluorescent channels, and the samples returned to a 37°C/5% CO_2_ incubator, with the samples re-imaged at 4, and 24 hours. All imaging was performed using identical excitation and camera settings to enable fluorescence quantification between timepoints. Imaging was performed on a Leca DMI6000B equipped with a Photometrics Evolve 512 EM-CCD camera, Sola light engine, Chroma Sedat Quad filter set with blue (Ex: 380/30, Em: 455/50), green (Ex: 490/20, Em: 525/36), red (Ex: 555/25, Em: 605/52), and far-red (Ex: 645/30, Em: 705/72) filter pairs, and the LAS-X software platform, using a 40×/1.30 NA objective lens. The resulting tiles were assembled into a single large image using LAS-X software, and individual macrophages segmented using the far-red channel and a trained machine learning algorithm in Ilastik [74]. The resulting cell masks were then imported into FIJI, and the integrated intensity of the Cell Proliferation Dye eFluor 670 in each cell quantified using the “Analyze Particles” tool [72].

### RT-PCR

Total RNA was isolated from monocytes, moMacs, and THP1 cells using a Qiagen RNA isolation kit and converted to cDNA using an iScript cDNA kit. q-PCR was run for samples using SsoFast Evagreen Supermix on a Quantico Q PCR thermocycler. Gene expression was calculated using the ΔΔCt method, using GAPDH as a reference gene. All primer sequences used in q-PCR are shown in **Supplemental Table 2**.

### Apoptosis Assays

THP1 macrophages were differentiated on glass coverslips as described above, and then stimulated for 24 hours with either 10 mmol/L oxLDL or 10 μM thapsigargin for 48 hr. The cells were then cooled to room temperature and stained with Alexa-488 conjugated AnnexinV in binding buffer (150 mM NaCl, 2 mM CaCl_2_, 20 mM HEPES, pH 7.4) for 10 min. The cells were washed once with binding buffer, fixed with 4% PFA in binding buffer, counter-stained with 1 μg/mL DAPI for 5 min, washed once with PBS, and then mounted on a glass slide using Permount Fluorescent Mounting Medium. An 11×11 tiled area of each sample was imaged as described in the proliferation assay, the resulting tiles were assembled into a single large image using LAS-X software, and the portion of cells positive for AnnexinV staining determined using the Cell Counting feature in FIJI [72].

### Statistics

Statistics were performed in Graphpad Prism version 9. Data is presented as individual measurements plotted over the mean ± SEM, and tested for normality using a Shapiro-Wilk test prior to analysis. Unless otherwise noted, all statistics were performed using non-parametric 2-tailed test, and with a significance cutoff of 0.05. The specific statistical tests used are indicated in the figure captions.

## Acknowledgements & Funding

We would like to thank the Molecular Imaging Facility (www.molimage.ca) and Kristin Chadwick of the London Regional Flow Cytometry Facility for their assistance with the microscopy and flow cytometry assays. This study was funded by a Heart and Stroke Foundation of Canada Grant-In-Aid to BH. AA was funded by an Ontario Graduate Scholarship and a Dr Robert George Everitt Murray Graduate Scholarship from The University of Western Ontario. PHC was funded via the X-Labs program at The University of Western Ontario. The funders had no role in study design, data collection and analysis, decision to publish, or preparation of the manuscript.

