## Supplemental Figures 1-5, Supplemental Tables 1-2 for "GATA2 Mediates Macrophage Proliferation During Atherosclerosis"

### Supplemental Materials

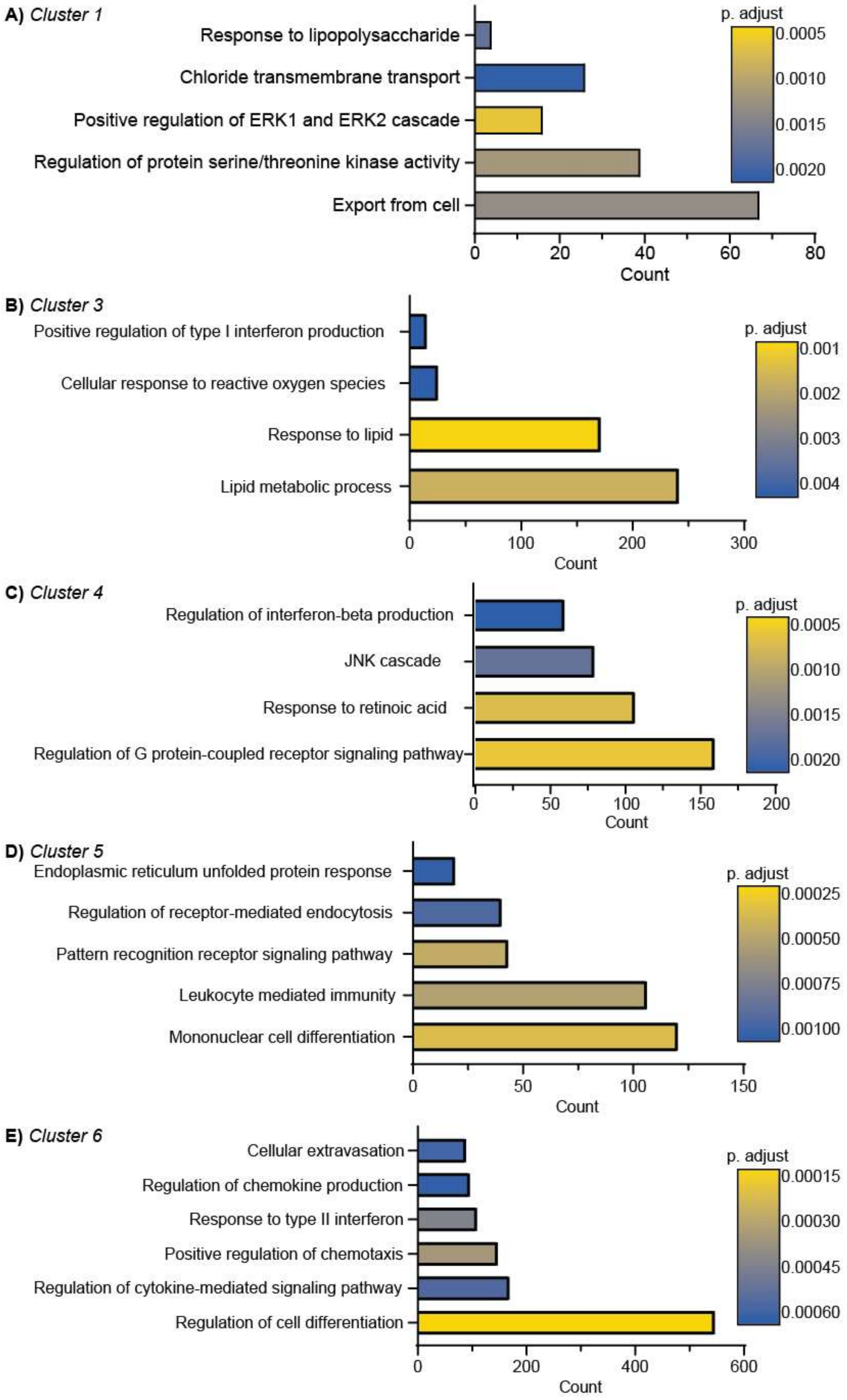

**Supplemental Figure 1: GO Analysis of Co-Regulated Gene Clusters.** Gene Ontology (GO) analysis of Cluster 1 (A) and Clusters 3-6 (B-E) of the transcriptomes of monocytes, GATA2+ macrophages, and monocyte-derived macrophages. Count is the number of genes in the cluster assigned to the corresponding GO annotation.

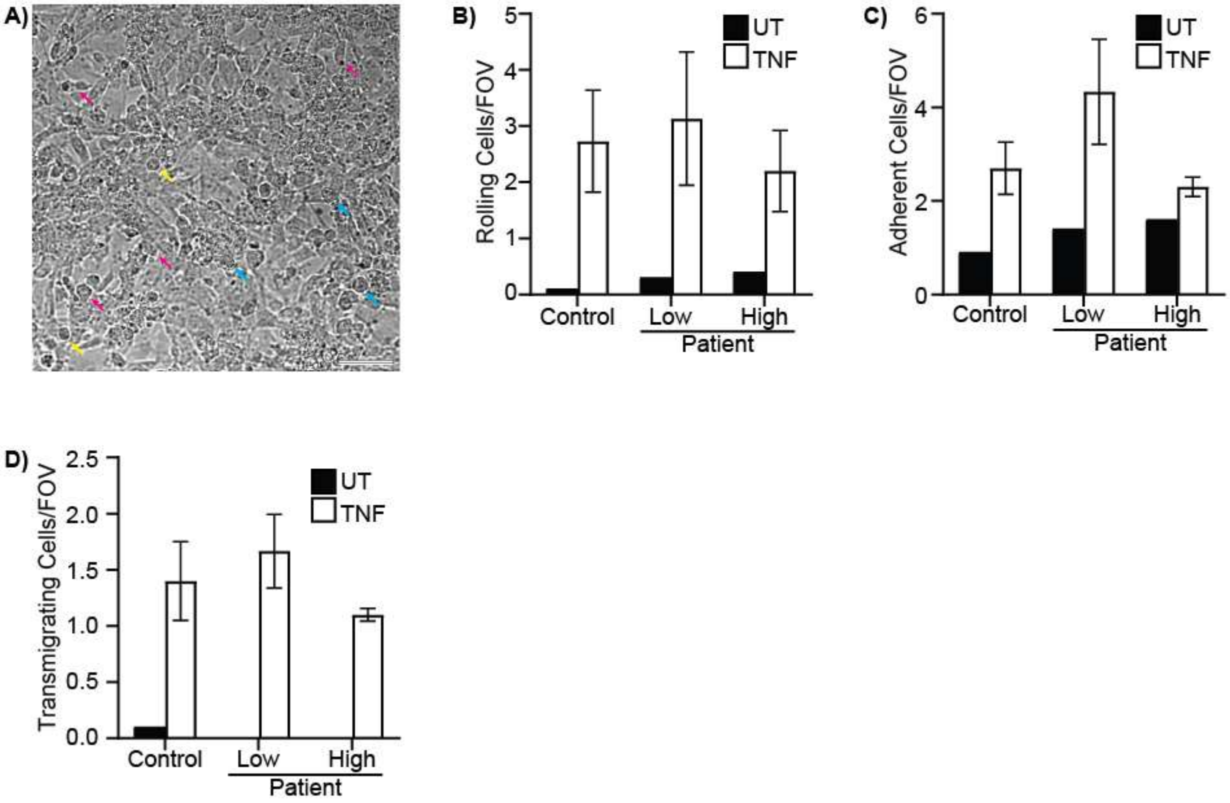


**Supplemental Figure 2: Monocyte Recruitment to Activated Endothelium.** Human aortic endothelial cells were cultured in a parallel plate flow chamber and stimulated with 5 ng/mL of TNFα for 4 hours, after which purified monocytes from healthy controls or atherosclerosis patients with low versus high monocytic GATA2 expression profuse at 50 dynes/$cm^2$. **A)** Image of monocytes from a high-GATA2 patient on TNFα-stimulated endothelium. Rolling cells are indicated with yellow arrows, adherent cells with cyan arrows, and transmigrating cells with magenta arrows. Scale bar is 50 µm. The number of rolling (**B**), adherent (**C**), and transmigrating (**D**) monocytes were quantified. No statistically significant differences ($p < 0.05$) were observed between untreated (UT) or TNFα-stimulated (TNF) endothelium, ANOVA with Tukey correction, n = 3.

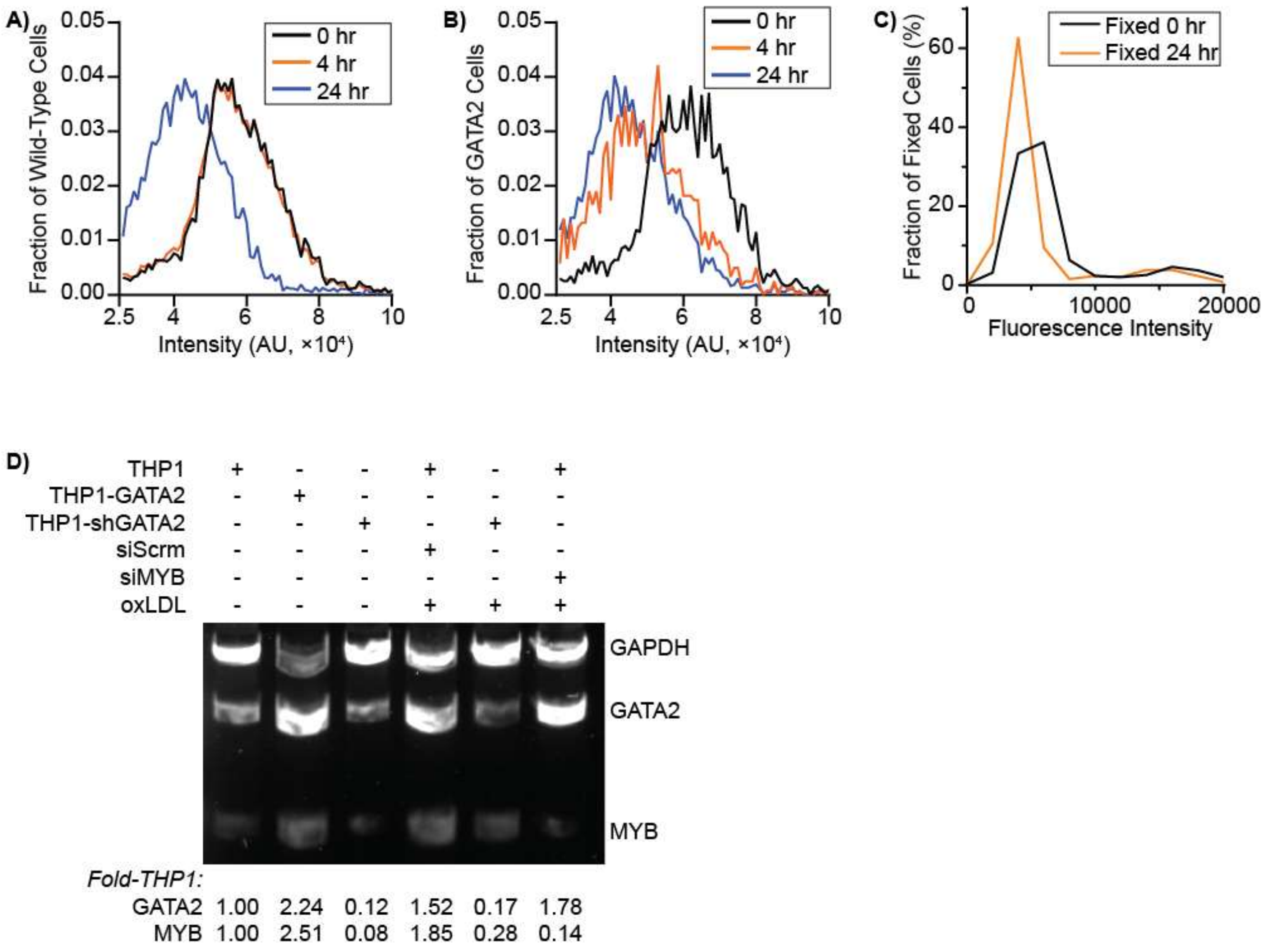


**Supplemental Figure 3: Macrophage Proliferation Assays.** THP1 cells were pre-stained with Cell Proliferation Dye eFluor 670 and differentiated into macrophages by addition of 100 ng/mL PMA, and the dilution of the Cell Proliferation Dye eFluor 670 used to quantify proliferation. **A,B)** Cell Proliferation Dye eFluor 670 intensity histograms in wild-type THP1 cells (A) and GATA2-overexpression THP1 cells (B) undergoing macrophage differentiation. Intensity data was collected at 0, 4, and 24 hours post-PMA stimulation. **C)** Cell Proliferation Dye eFluor 670 histogram for wild-type THP1 cells that were fixed with 4% paraformaldehyde immediately after labeling. **D)** Quantification of siRNA and GATA2 overexpression efficacy by semi-quantitative qPCR. GATA2 and MYB expression were measured in wild-type THP1 cells (THP1) or in THP1 cells over-expressing GATA2 (THP1-GATA2) or expressing a GATA2-targeting shRNA (THP1-shGATA2). In some experiments, cells were pre-treated for 24 hours with 2.5 mmol/L oxLDL (oxLDL), following a 48-hour treatment with either a scrambled (siScrm) or MYB-targeting (siMYB) cell-penetrant siRNA. Numbers below the blot are densitometry of the GATA2 and MYB bands, normalized to the intensity of GAPDH in the same sample, then normalized to untreated THP1 cells (first lane). Data is representative of a minimum of 3 experiments.

**Supplemental Table 1: Antibodies and Dyes Used in this Study**

| Name of Antibody or Dye | Conjugated Fluorophore | Company | Catalog | Clone | Dilution |
|---|---|---|---|---|---|
| Fixable viability dye | eFluor 780 | Thermo-Fisher | 65-0865-14 | | 1:1000 |
| Hoechst 33342 | | Thermo-Fisher | 62249 | | 1 μg/mL |
| Cell Proliferation Dye eFluor 670 | eFluor 670 | Thermo-Fisher | 65-0840-85 | | 10 μM |
| CD14 | APC | BioLegend | 367117 | 63D3 | 1:100 |
| GATA2 | Alexa Fluor 555 | Abcam | ab109241 | EPR2822 | 1:100 |
| Ki-67 | FITC | BioLegend | 350532 | Ki67 | 1:100 |
| Ki-67 | Unconjugated | ThermoFisher Scientific | MA5-14520 | SP6 | 1:100 |
| CD68 | Unconjugated | Abcam | ab201340 | C68/684 | 1:100 |
| Ki-67 | Alexa Fluor 350 | R&D systems | IC7617U-100UG | 1297A | 1:100 |

**Supplemental Table 2: Primers Used in this Study**

| Name of Gene | Primer Sequence | Tm (°C) |
|---|---|---|
| GAPDH forward primer | 5'-TCAAGGCTGAGAACGGGAAG-3' | 57 |
| GAPDH reverse primer | 5'-CGCCCCACTTGATTTTGGAG-3' | 56.7 |
| GATA2 forward primer | 5'-CAGCAAGGCTCGTTCCTGTTCA-3' | 59.9 |
| GATA2 reverse primer | 5'-ATGAGTGGTCGGTTCTGCCCAT-3' | 60.9 |
| Ki-67 forward primer | 5’-TCCTTTGGTGGGCACCTAAGACCTG-3’ | 62.5 |
| Ki-67 reverse primer | 5’-TGATGGTTGAGGTCGTTCCTTGATG-3’ | 58.9 |
| CCR2 forward primer | 5’-CAGGTGACAGAGACTCTTGGGA-3’ | 58.1 |
| CCR2 reverse primer | 5’-GGCAATCCTACAGCCAAGAGCT-3’ | 59.5 |
| Integrin $\alpha_4$ forward primer | 5’-GCATACAGGTGTCCAGCAGAGA-3’ | 58.8 |
| Integrin $\alpha_4$ reverse primer | 5’-AGGACCAAGGTGGTAAGCAGCT-3’ | 60.6 |
| Integrin $\beta_2$ forward primer | 5’-AGTCACCTACGACTCCTTCTGC-3’ | 58.2 |
| Integrin $\beta_2$ reverse primer | 5’-CAAACGACTGCTCCTGGATGCA-3’ | 59.9 |
| MYB forward primer | 5’-GAAAGCGTCACTTGGGGAAA-3’ | 61.1 |
| MYB reverse primer | 5’-TGTTCGATTCGGGAGATAATTGG-3’ | 60.2 |
| JUN forward primer | 5’-GCCAGGTTCAGGGTCATGC-‘3 | 61.1 |
| JUN reverse primer | 5’-AAACTAACCTCACGTGAAGTGACG-3’ | 61.4 |
